# Sparse coding and dimensionality reduction in cortex

**DOI:** 10.1101/149880

**Authors:** Michael Beyeler, Emily Rounds, Kristofor D. Carlson, Nikil Dutt, Jeffrey L. Krichmar

**Affiliations:** University of California, Irvine, Irvine, CA, USA; University of Washington, Seattle, WA, USA; Sandia National Laboratories**, Albuquerque, NM, USA

**Keywords:** efficient coding, sparse coding, dimensionality reduction, nonnegative matrix factorization, neural computation, spike-timing dependent plasticity

## Abstract

Supported by recent computational studies, sparse coding and dimensionality reduction are emerging as a ubiquitous coding strategy across brain regions and modalities, allowing neurons to achieve nonnegative sparse coding (NSC) by efficiently encoding high-dimensional stimulus spaces using a sparse and parts-based population code. Reducing the dimensionality of complex, multimodal sensory streams is critically important for metabolically constrained brain areas to represent the world. In this article, we provide an overview of NSC, summarize evidence for its role in neural computation in disparate regions of the brain, ranging from visual processing to spatial navigation, and speculate that specific forms of synaptic plasticity and homeostatic modulation may underlie its implementation. We suggest that NSC may be an organizing principle in the nervous system.

## A need for compressed representations of a high-dimensional world

Brains face the fundamental challenge of processing, storing, and representing high-dimensional stimuli using patterns of neural activity. This challenge is complicated by strict constraints on metabolic cost [1] and the widespread existence of anatomical bottlenecks [2-5], which often force the information stored in a large number of neurons to be compressed into an orders of magnitude smaller population of downstream neurons; for example, storing information from 100 million photoreceptors in 1 million optic nerve fibers, or resulting in a 10 – 10, 000 fold convergence from cortex to the basal ganglia [5].

One potential approach to addressing this challenge is to reduce the number of variables required to represent a particular stimulus space; a process known as **dimensionality reduction** (see Glossary). This idea features prominently in efficient coding theories of brain function [6-9], which posit that the brain performs dimensionality reduction by maximizing mutual information between the high-dimensional input and the low-dimensional output of neuronal populations. Several modern variants of the efficient coding hypothesis, such as independent component analysis (ICA) [10] and nonnegative sparse coding (NSC) [11, 12], suggest that the role of cortical representations is to further reduce the redundancy of the sensory signal by separating it into its underlying causes, a process known as **factor analysis**.

Here we propose that a variety of neuronal responses can be understood as an emergent property of efficient population coding based on dimensionality reduction and sparse coding. Specifically, computational evidence from data analyses and computer simulations suggest that NSC, a combination of nonnegative matrix factorization (NMF) [13, 14] and sparse population coding [15] (Box 1), generates low-dimensional embeddings of high-dimensional stimulus spaces that resemble neuronal population responses in a variety of brain regions. Reminiscent of **basis function** representations [16-18], these embeddings are both sparse and parts-based, allowing for perceptual or behavioral variables of interest to be represented by taking a linear combination of neuronal responses. Furthermore, we suggest that these computations might be implemented in the brain through mechanisms of synaptic plasticity and homeostasis, which were previously shown to be capable of performing statistical inference on synaptic inputs [19-22].

Overall, our observations suggest that commonly reported neuronal response properties might simply be a by-product of neurons performing a biological equivalent of dimensionality reduction on their inputs via Hebbian-like learning mechanisms.

## Nonnegative sparse coding as a modern variant of the efficient coding hypothesis

Early theories of efficient coding [6, 23] were developed based on the visual system. Realizing that the set of natural scenes was much smaller than the set of all possible images, it was argued that the visual system should not waste resources on processing arbitrary images. Instead, the nervous system should use statistical knowledge about its environment to represent the relevant input space as economically as possible.

Modern renditions of this theory (such as sparse coding [24] and ICA [10]) have refined the original hypotheses by tying neuronal response properties to the statistics of the natural environment (for a review, see [25]). These theories were shaped by two fundamental empirical observations about early visual cortex: 1) a neuron’s **receptive field (RF)** resembled a decomposition of the visual stimulus into a series of local, largely independent feature components (e.g., a 2-D Gabor function is basically a local approximation of the directional spatial derivative of an image), and 2) any individual neuron responded only sparsely to a small subset of stimulus features (e.g., orientation or color at a particular spatial location). Thus a neuron’s RF could be understood as a sparse, low-dimensional embedding of high-dimensional input stimuli. Such a representation is desirable from an energy-expenditure point of view, since it allows the visual system to represent any visual stimulus by activating only a small set of neurons, while most neurons in the population remain silent.

Olshausen and Field [24] found that linear sparse coding of natural images yielded features qualitatively similar to RFs of simple cells in primary visual cortex (V1), thus giving empirically observed RFs an information-theoretic explanation. However, as pointed out by Hoyer [26], sparse coding falls short of providing a literal interpretation for V1 simple-cell behavior for two reasons: 1) every neuron could be either positively or negatively active, and 2) the input to the neural network was typically double-signed, whereas V1 neurons receive visual input from the lateral geniculate nucleus (LGN) in the form of separated, nonnegative ON and OFF channels.

Hoyer [11, 26] proposed NSC as a way to transform Olshausen and Field’s sparse coding from a relatively abstract model of image representation into a biologically plausible model of early visual cortex processing by enforcing both input signal and neuronal activation to be nonnegative. This seemingly simple change had remarkable consequences on the quality of the sensory representation: Whereas elementary image features in the standard sparse coding model could “cancel each other out” through subtractive interactions, enforcing nonnegativity ensured that features combined additively, much like the intuitive notion of combining parts to form a whole. The resulting *parts-based* representations resembled RFs in V1 more closely than other *holistic* representations, and would later be shown to apply to other brain regions as well, suggesting that parts-based representations are employed throughout cortex.

Inhibitory connections can be modeled in the same fashion, by interpreting inhibitory weights as nonnegative synaptic conductances, which not only preserves the parts-based quality of the encoding, but also allows for more complicated connection types to be modeled (e.g., V1 neurons receiving input from both excitatory ON and inhibitory OFF cells in the LGN) [26]. However, it is interesting to note that a more recent study has argued that the nonnegativity constraint on the synaptic weights might not be necessary to preserve the parts-based quality of the encoding [27].

## Empirical evidence for nonnegative sparse coding in the brain

### Reconstructing stimulus spaces using sparse, parts-based representations

An influential paper by Lee and Seung [14] found that applying a linear dimensionality reduction technique known as NMF to a database of face images yielded sparse, localized features that resembled parts of a face. NMF is a statistical matrix decomposition technique that takes an *F* × *S* data matrix **V** whose rows correspond to distinct features of the input (e.g., *F* different pixels of an image) and whose columns correspond to different stimuli or observations of those features (e.g., *S* different images). Matrix **V** must be nonnegative, and NMF decomposes the matrix into two reduced-rank matrices whose linear combination can be weighted such that the product of **W** and **H** provides an accurate reconstruction of **V** (Fig. 1).

**Figure 1:**
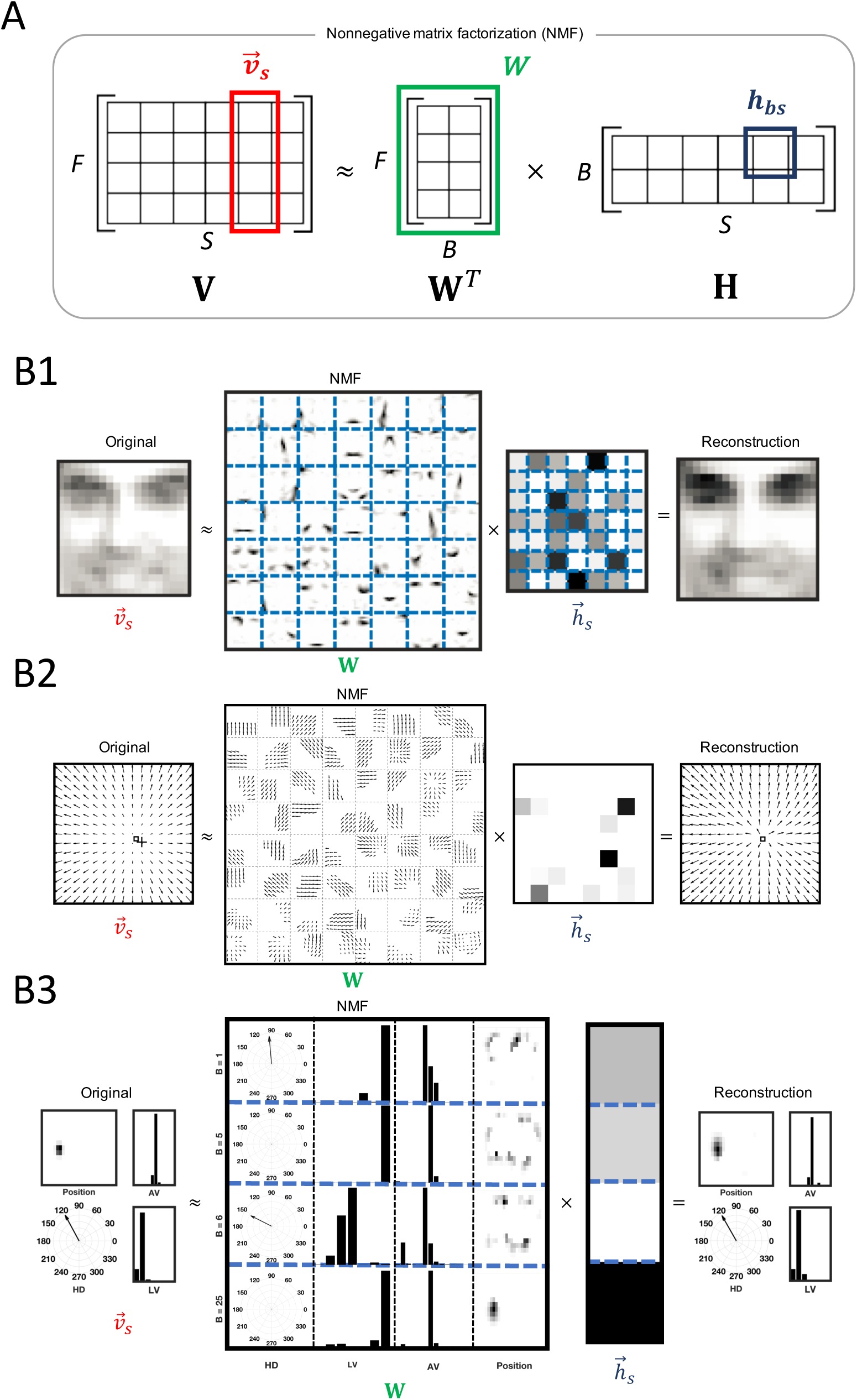
(Key Figure) NMF can recover sparse, parts-based representations of high-dimensional input data. A) A data matrix **V** (*F* features x *S* stimulus observations) can be reconstructed from two reduced-rank matrices **W** (containing *B* basis vectors) and **H** (containing the hidden coefficients of the decomposition). Any individual input stimulus (i.e., column in **V**) can be reconstructed from a linear combination of a set of basis vectors (i.e., the columns in **W**). B1-B3) Recovered weight matrices **W** resemble the receptive fields of cortical neurons across brain regions. The contribution of each basis vectors is given by a column in **H** (the darker the color, the stronger the contribution). B1) A facial image can be reconstructed from a sparse activation of simulated neurons in inferotemporal cortex that preferentially respond to parts of faces (adapted from [14]). B2) An optic flow field can be reconstructed from a sparse activation of MSTd neurons that prefer various directions of 3D self-translation and self-rotation (adapted from [31]). B3) A rat’s allocentric position and route-based direction of motion can be reconstructed from a sparse activation of model retrosplenial cortex (RSC) neurons that prefer an intricate combination of linear velocity (LV), angular velocity (AV), head direction (HD) and position (Position). Only the 4 most contributing hidden coefficients (those whose coefficients exceeded the mean of all coefficients) out of 30 are shown for the sake of clarity.

On its surface, NMF would appear to be unrelated to the mechanisms underlying artificial or biological neural networks; however, it was Lee and Seung’s intuitive mapping of these variables onto a neural network that forms the cornerstone of our conception of the NSC framework: A particular image, in this case encoded by *F* = 19 × 19 = 361 pixels *v*_1_, … , *v_F_* (i.e., a column of **V**), could be reconstructed by a linear mixing of a total of *B* encoding variables *h*_1_, … , *h_B_* (i.e., a column of **H**). A single encoding variable *b* ∊ 1, … , *B* influences multiple image pixels, owing to the fan-out of the connections from the encoding variable. As a result, a particular image of a face (Fig. 1B1) could be accurately represented by a linear combination of a small number (*B* = 49) of encoding variables or “basis images”. Such a representation is reminiscent to neural processing in inferotemporal cortex (IT), an area in the ventral visual “what” stream involved in encoding high-level object identity [28, 29], where images of whole faces can be linearly reconstructed using responses of approximately 200 neurons that each respond to certain sets of physical facial features [30].

Remarkably, such a parts-based representation is not specific to information processing in IT; the same principle can be extended to other areas of the visual system, such as the dorsal subregion of the medial superior temporal area (MSTd), which is part of the visual motion pathway [31]. Neurons in MSTd respond to relatively large and complex patterns of retinal motion (“optic flow”), owing to input from direction and speed selective neurons in the middle temporal area (MT) (for a recent review, see [32]). Although MSTd had long been suspected to be involved in the analysis of self-motion, the complexity of neuronal response properties has made it difficult to experimentally investigate how neurons in MSTd might perform this function. However, when Beyeler and colleagues [31] applied NMF to MT-like patterns of activity, they found a sparse, parts-based representation of retinal flow (Fig. 1B2) similar to the parts-based representation of faces encountered by Lee and Seung [14]. The resulting “basis flow fields” showed a remarkable resemblance to receptive fields of MSTd neurons, as they preferred an intricate mixture of 3D translational and rotational flow components in a subset of the visual field. As a result, any flow field possibly to be encountered during self-movement through a 3D environment could be represented by only *B* = 64 MSTd neurons, as compared to *F* = 9, 000 simulated MT input neurons. This led to an efficient and sparse population code, where any given stimulus could be represented by only a small number of simulated MSTd neurons (population sparsity), and any given model neuron was activated by only a small number of stimuli (lifetime sparsity) [31].

Analogously, NSC can explain response properties of neurons outside the visual system, such as in the retrosplenial cortex (RSC), an area important for navigation and spatial memory [33-35]. Neurons in the RSC conjunctively encode multiple variables related to the environment and one’s position and movement within it, allowing the representation of spatial features of the environment with respect to multiple reference frames [36], as determined by an experiment in which rats ran back and forth on a W-shaped track that occupied two different spatial locations within the room in each recording session to test sensitivity to the allocentric reference frame (i.e., track positions *α* and *β*). Outbound and inbound runs were made up of opposite turn sequences (left-right-left (LRL) and right-left-right (RLR), respectively) that corresponded to different sets of trials, which allowed assessment of sensitivity to the egocentric and route-based reference frames. However, establishing a mechanistic link between physiological response properties of RSC neurons and their underlying representations of space has proved difficult, due to the complexity of their response properties and because inputs to the region are not easily isolated.

We applied NMF to recorded data from the original RSC experiment conducted by Alexander and Nitz [36]. During the experiment, activity from 228 RSC neurons was recorded along with four behavioral metrics: linear velocity, angular velocity, head direction, and allocentric position. Using Gaussian and cosine tuning curves, we created idealized input neurons that encoded these four variables. By applying NMF to the idealized neuronal output, we were able to replicate functionality observed in the biological RSC (Fig. 1B3). Once again, the dimensionality was reduced from a set of *F* = 417 input neurons to a set of *B* = 30 basis functions.

Although there seems to be a consensus that information-theoretic explanations are relevant when investigating the early visual system, higher-order brain areas are often considered to be specialized for performing tasks (e.g., recognizing objects, making decisions, navigating an environment), rather than the efficient encoding of information. However, the finding that NSC could be used to explain neuronal responses in the visual and retrosplenial cortices introduces the possibility that it might apply elsewhere in the brain. In fact, sparse (and potentially parts-based) representations have been observed in olfactory, auditory, somatosensory, and motor cortices (Table 1). This introduces the possibility that NSC might in fact be a general principle to which neuronal computations adhere.

**Table 1:** Nonnegative sparse coding in the brain. We list a group of brain regions for which there is experimental evidence of certain features associated with NSC (‘X’: evidence exists, ‘?’: has yet to be investigated). For each brain region, the left-hand side of the table lists experimental evidence for sparse and/or parts-based representations, whereas the right-hand side lists computational support that NSC can describe receptive fields or response properties within that region.

| **Area** | **Sparse** | **Parts-**<br>**based** | **Experimental**<br>**evidence** | **Modeled**<br>**by NSC** | **Computational**<br>**support** |
| --- | --- | --- | --- | --- | --- |
| Retina | X | X | [27, 37] | X | [27, 37] |
| Early visual cortex | X | X | [24, 26, 38, 39] | X | [20, 24, 26, 40] |
| Ventral visual stream | X | X | [30, 41, 42] | X | [14, 43] |
| Dorsal visual stream | X | X | [16, 17, 44] | X | [31] |
| Auditory cortex | X | ? | [45] | ? | ? |
| Olfactory cortex | X | ? | [46] | ? | [21] |
| Retrosplenial cortex | X | X | [36] | X | present paper |
| Motor cortex | X | ? | [47, 48] | ? | [49] |
| Basal ganglia | X | ? | [3, 50] | X | [3], advanced RDDR |
| Barrel cortex | X | ? | [51] | ? | ? |
| Hippocampus | X | ? | [52] | ? | ? |

### Understanding neuronal response properties within the framework of NSC

Neuronal response properties can also be computed for model neurons with NSC, which means that model neurons can be evaluated using methods similar to the ones employed by experimental researchers to understand biological neurons, and by theoretical neuroscientists to understand computational models. This is important because it means that NSC can be used to model neural activity in the brain, and the resulting activity patterns generated by NSC can be compared to and evaluated against experimental findings.

Neuronal response patterns can be computed by reconstructing the activity of a population of “input” neurons (i.e., the *s*-th column in **V**) from a population of “output” neurons (i.e., the *s*-th column in **H**) via their synaptic weights (i.e., the matrix **W**; Fig. 1A). Similarly, it is also possible to calculate **H** from **W^T^** and **V** (Fig. 2A). In this context, a single element of **H** corresponds to the activity of the *b*-th neuron to the *s*-th stimulus, which is given by the dot product of its presynaptic connections (i.e., the *s*-th column in **V**) and the corresponding synaptic weights (i.e., the *b*-th row in **W^T^**). Note that the response of the model neuron to different stimuli *s* ∊ 1, … , *S* involves different columns of **H** and **V**, but always relies on the same weight matrix **W**. Thus, we can utilize **W** (which must remain fixed once learned with NMF) to simulate a model neuron’s response to arbitrary input stimuli by replacing the column in **V** with new input. This allows us to investigate the response properties of individual model neurons much in the same way that experimental neuroscientists study biological neurons.

**Figure 2:**
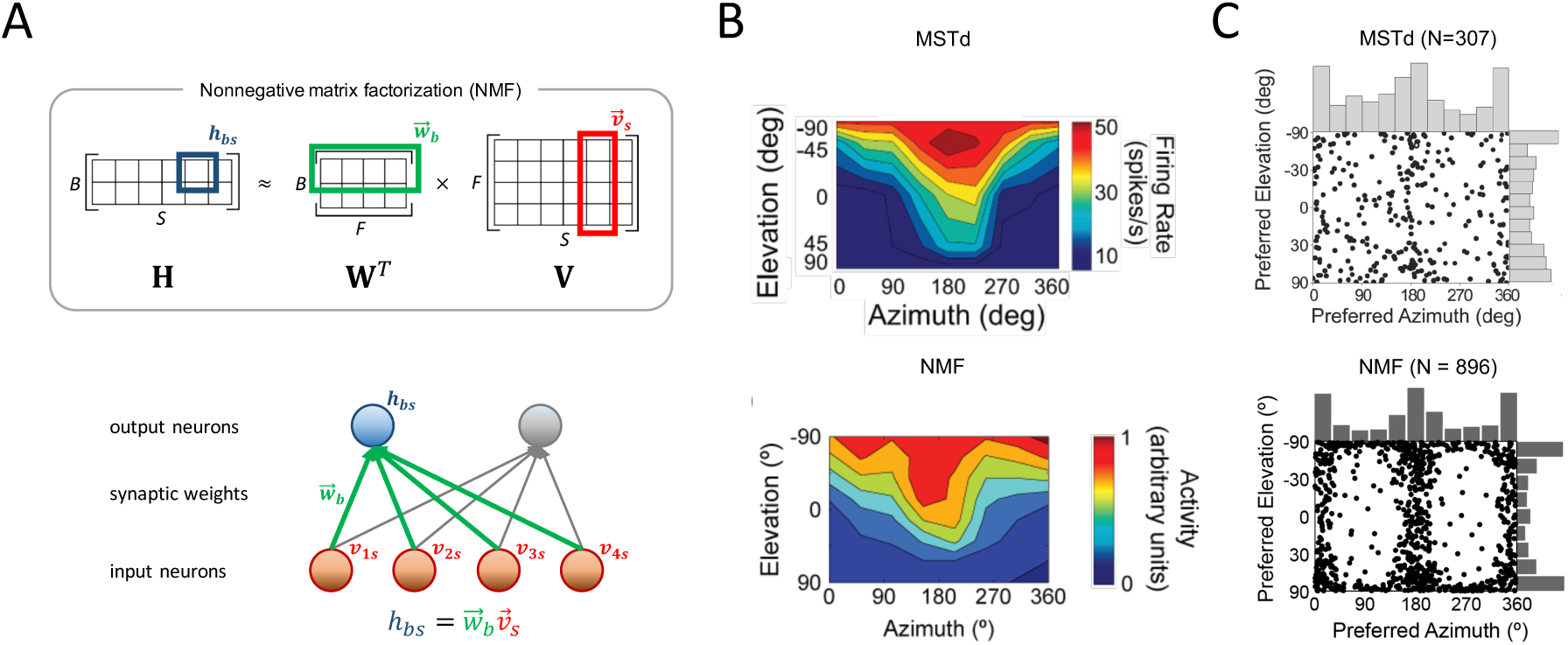
NSC can predict response properties of biological neurons. A) Solving the NMF equation for **H**, the response of output neuron *b* to stimulus *s* (i.e., matrix element *h_bs_*) can be calculated from the activity of all input neurons (i.e., column *s* in **V**) and the corresponding synaptic weights (i.e., row *b* in **W***^T^*). B) Example of 3D heading tuning for a neuron in macaque MSTd (top, reprinted from [53]) and a simulated neuron (bottom, reprinted from [31]). Color contour maps show the mean firing rate or model activation as a function of azimuth and elevation angles of the self-movement direction in 3D. C) Classifying simulated akin to biological neuronal responses yields distributions of 3D heading preferences akin to macaque MSTd (top, reprinted from [31]). Each neuron also responded to a preferred direction of 3D self-rotation (not shown).

For example, subjecting model MSTd neurons to simulated optic flow fields mimicking natural viewing conditions during locomotion, Beyeler and colleagues [31] found that individual model neurons preferentially responded to a particular 3D direction of self-translation and self-rotation, much like individual neurons in macaque MSTd (Fig. 2B). In addition, known statistical properties of the MSTd population emerged naturally from the NMF based model, such as a relative over-representation of lateral headings (Fig. 2C).

Other groups have successfully applied methods similar to NSC to model neuronal response properties. For example, in the visual system, an NMF-based model was able to reconstruct neuronal spike trains in the salamander retina [37]. Following testing on ground truth data, the researchers recorded spikes from in vitro retinal ganglion cells while the cells were exposed to natural scenes (either still photographs or videos). They then applied several factorization methods to the data. Space-by-time NMF could decompose the data into separate spatial and temporal modules that yielded sparser and more compact representations compared to other techniques, including orthogonal Tucker-2 and basic NMF. NSC could be used to reproduce response properties of V1 simple and complex cells [26, 38] as well as V2 hypercomplex cells [54]. Outside the visual stream, a model known as Reinforcement-Driven Dimensionality Reduction (RDDR) successfully used Hebbian learning to reproduce basal ganglia response patterns associated with reward [50], a function associated with cortico-striato-pallidal circuitry. The authors later applied nonnegativity constraints to the Hebbian learning in the model so that it performed NMF on its inputs. The model advanced understanding of the cortico-striato-pallidal loop by capturing behavior of the circuit while explaining the existence of convergent and lateral connections in the region that other models have historically ignored [3]. The authors suggest that the basal ganglia uses unsupervised, reward-driven learning to perform dimensionality reduction on cortical inputs for the efficient compression of information in order to plan upcoming actions in the frontal cortex.

The finding that NSC can account for neuronal response properties across brain areas and modalities supports the view that these preferences emerge from the pressure to find efficient representations of perceptually or behaviorally relevant stimulus spaces. At the population level, NSC promotes representations in which neurons act as generalists rather than specialists, allowing for the simultaneous encoding of multiple variables of interest (e.g., heading, eye rotation velocity in MSTd [31]) with respect to multiple frames of reference (e.g., egocentric, route-based in RSC [55]). Among the advantages of such basis function representations [16-18] (also called mixed-selectivity representations [56-58]) are robustness to noise as well as the ability to decode various variables of interest through a linear combination of neuronal responses.

### Approximating NMF with STDPH

That NSC can explain and reproduce response properties observed in biological neurons may be an important clue as to how brains have evolved to parse and store information. One candidate NSC process in the brain is synaptic plasticity which, similar to NSC, may act on synapses for the purpose of reducing dimensionality on an input space in order to represent it efficiently and sparsely. Specifically, we propose that NSC might be functionally equivalent to spike-timing dependent plasticity and homeostatic synaptic scaling (STDPH) (Box 2). **Spike-timing dependent plasticity (STDP)** is a spike-timing based correlative learning rule in which the timing of the preand post-synaptic spikes determine the amplitude and sign of synaptic weight changes (i.e., long-term depression or long-term potentiation) [59, 60]. Homeostasis acts to keep neuronal activity in a good working range, acting across synapses of a single neuron and across multiple neurons [61]. Whereas STDP acts over a relatively short time scale (i.e., milliseconds to seconds), homeostasis operates over minutes to days. Homeostasis also facilitates synaptic competition by normalizing the inputs to a neuron. This evens the playing field for synapses that weaken due to imbalances in activity, but might otherwise strengthen if left to their own devices [62].

Experimental and theoretical evidence suggest that biological processes, such as synaptic plasticity and homeostatic mechanisms, may be reducing the dimensionality of inputs in a similar way to NMF. For example, Carlson and colleagues [20] delivered a mathematical proof that STDPH can approximate the NMF algorithm (see Box 2). Similar to Oja’s rule [22], which was developed to stabilize rate-based Hebbian learning (effectively resulting in principal component analysis (PCA)), synaptic scaling acts as a homeostatic mechanism to stabilize STDP (effectively resulting in NMF). This finding suggests that neurons are able to find accurate factorial representations of their input stimulus spaces through means of Hebbian-like synaptic learning. In addition, sparsity of the encoding might be enforced by spike thresholding [63] and lateral inhibitory connections [64].

To investigate this equivalence in a more realistic setting, we conducted simulations in which a model constructed using STDPH and a model constructed using NSC were applied to a dataset of recorded neuronal activity observed in the RSC of rats in a spatial navigation task (as shown for NSC in Figure 1B3). In the STDPH experiment, the idealized input neurons generated spike trains as input to a population of SNNs that were trained using an evolutionary strategy (Box 3) to match the recorded electrophysiological data [55]. In the NSC experiment, the idealized neuronal output was used to construct a matrix of training data associated with each of the four recorded ‘features’ (linear velocity, angular velocity, head direction, and position), to which NMF was applied.

We found that the activity patterns of both NSC and STDPH model neurons could replicate the neuronal response properties and ensemble activity seen in the electrophysiologically recorded neurons in the dataset (Fig. 3A, B, C, left); that is, the model neuron activity could be classified into three broad categories, with remarkably similar population statistics to rat RSC [36]: 1) Turn-sensitive, no modulation neurons responded whenever the animal made a left or right turn on the track (light gray); 2) Turn-sensitive, route modulation neurons responded whenever the animal made a turn on a specific position along the route, independent of allocentric location (dark gray); and 3) Turn-insensitive neurons that could not be classified according to the above, but nonetheless exhibited complex and robust firing patterns (white). In addition, both NSC and STDPH model RSC produced simulated neurons whose ensemble activity vectors could be used to predict the agent’s location on a route with respect to the allocentric frame of reference. Ensemble activity patterns could also disambiguate the agent’s location within routes that occupied different locations in the room, consistent with findings of population behavior in the biological RSC [36]. When even and odd trials on the same track locations were compared, ensemble prediction error was very low, but when the tracks were in two different locations (i.e., *α* vs. *β*), the prediction error was significantly higher in all cases (Fig. 3A, B, C, right). For further details on this analysis, see [36, 55].

**Figure 3:**
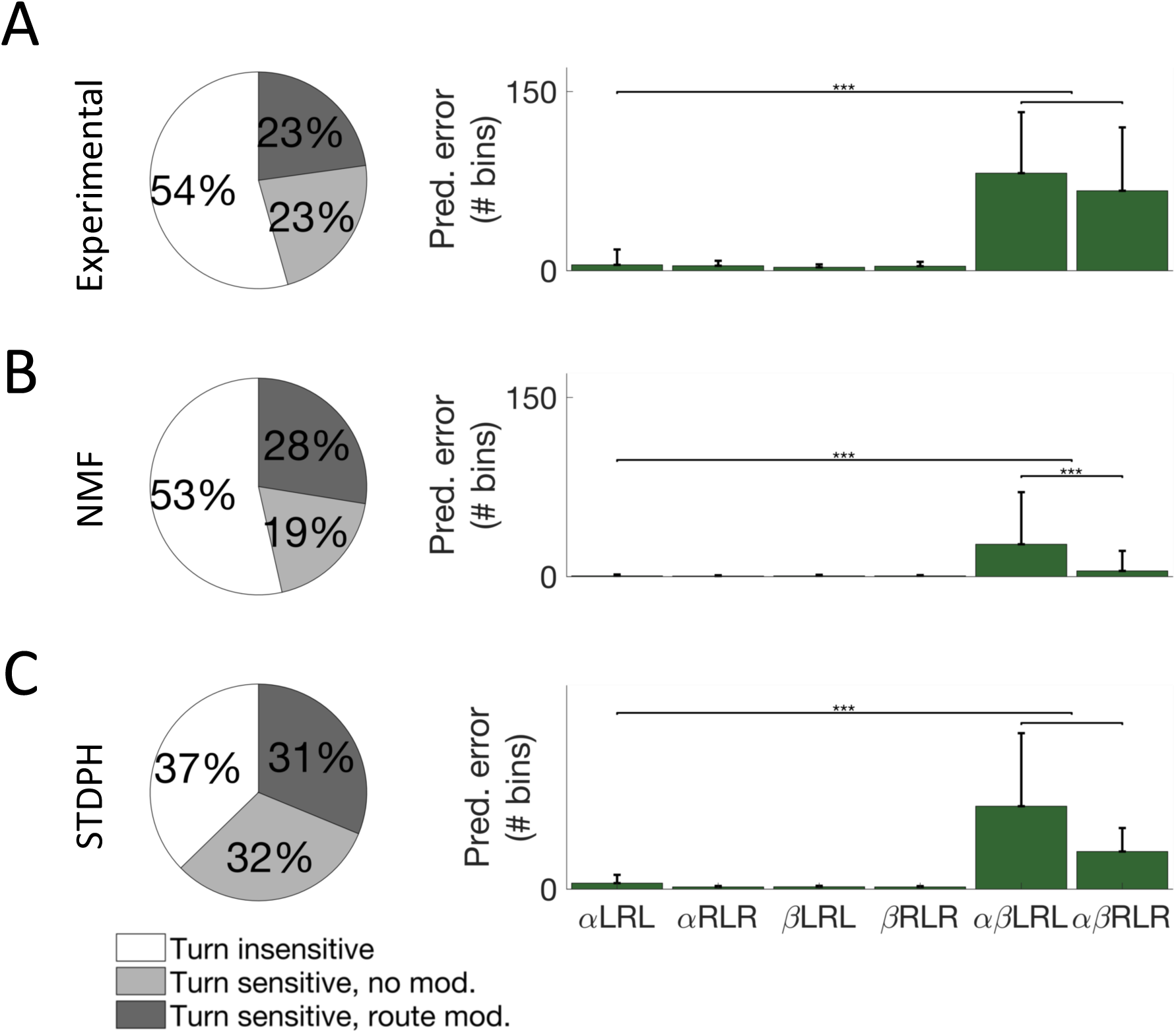
Functional neuron type distributions found in the dataset and produced by each model (left columns) and average error corresponding to positional ensemble reconstruction matrices (right columns). Separate allocentric positions of the track are represented by the symbols *α* and *β*. Reconstructions between the track positions are represented by *αβ*. There were two possible routes associated with each track: an outbound run consisting of a left-right-left (LRL) turn sequence and an inbound run consisting of a right-left-right (RLR) turn sequence. A) Experimental data from [36]. B) Simulated using NMF. C) Simulated by evolving STDPH parameters to fit experimental data from [55].

These findings lend credence to our proposal that NMF and STDPH are functionally equivalent. Furthermore, the matrix **W** produced by NMF when applied to the RSC dataset was qualitatively similar to the synaptic weight matrices produced by simulations in which STDPH was evolved to match the same dataset.

## Concluding remarks and future directions

In this article, we have argued that a variety of neuronal response properties can be understood as an emergent property of efficient population coding based on dimensionality reduction and sparse coding. We offer three testable predictions of this theory:

First, we predict that parts-based representations can explain RFs of neurons in a variety of brain regions, including but not limited to those brain areas discussed here (i.e., V1, MSTd and RSC). In agreement with the literature on basis function representations [16-18], we expect parts-based representations to be prevalent in regions where neurons exhibit a range of tuning behaviors [31], display mixed selectivity [56, 57], or encode information in multiple reference frames [36, 55].

Second, where such representations occur, we expect the resulting neuronal population activity to be sparse, in order to encode information both accurately and efficiently. Sparse codes offer a trade-off between dense codes (where every neuron is involved in every context, leading to great memory capacity but suffering from cross talk among neurons) and local codes (where there is no interference, but also no capacity for generalization) [65].

Third, we propose that STDPH is carrying out a similar function to NMF, and may be attempting to approximate the linear RF of neurons participating in sparse, parts-based representations throughout the brain. STDPH and NMF can effectively produce NSC. With the emergence of computational tools developed to understand the neural code in high stimulus dimensions [66], we expect to see qualitative similarities between empirically observed RFs and those recovered by NMF and STDPH. Such findings would be consistent with the idea that neurons can perform statistical inference on their inputs via Hebbian-like learning mechanisms [19-22].

In summary, we suggest that the evidence reviewed in this paper may indicate that dimensionality reduction through nonnegative sparse coding, implemented by synaptic plasticity rules, may be a canonical computation throughout the brain.

### Box 1: Nonnegative sparse coding (NSC)

Nonnegative sparse coding (NSC) combines nonnegative matrix factorization (NMF), a linear dimensionality reduction technique from statistical learning, with sparse population coding from neural network theory [11, 67]. NMF belongs to a class of methods that can be used to decompose a multivariate data matrix **V** into an inner product of two reduced-rand matrices **W** and **H**. NMF assumes that the observed data in **V** are caused by a collection of latent factors weighted by nonnegative numbers, representing both the presence and the intensity of the cause.

In the context of NSC, **V** and **H** correspond to activation values of two distinct neuronal populations, which are connected to each other via synaptic weight values in **W** (Fig. 1). Consider a number of data samples *s* ∊ [1, *S*], for example, in the form of observed firing rates of a population of *F* neurons. If we arrange the observed values of the *s*-th observation into a vector *v_s_*, and if we arrange all vectors into the columns of a data matrix **V**, then linear decompositions describe these data as **V** ≈ **WH**. Here, **W** is a matrix that contains as its columns a total of *B basis vectors* of the decomposition, and **H** contains as its rows the *hidden coefficients* that give the contribution of each basis vector in the input vectors. The difference between **V** and **WH** is termed the *reconstruction error*.

The goal of NSC is then to find a linear decomposition of **V** that minimizes the reconstruction error, while guaranteeing that both **W** and **H** are sparse. This can be achieved by minimizing the following cost function [11]:
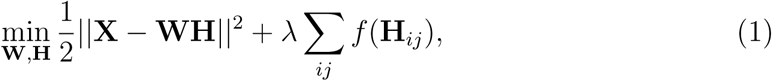

subject to the constraints ∀*ij* : **W***_ij_* ≥ 0, **H***_ij_* ≥ 0, and 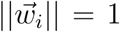, where 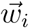 denotes the *i*th column of **W**. Here, the left-hand term describes the reconstruction error, which can be minimized with NMF, and the right-hand term describes the sparseness of the decomposition. The trade-off between sparseness and accurate reconstruction is controlled by the parameter λ (λ ≥ 0), whereas the form of *f* defines how sparseness is measured (a typical choice is the L1 norm on **H**).

Note that several modern implementations of the NMF algorithm, such as Matlab’s `nnmf.m` function, already implement sparsity constraints, hence making NMF indistinguishable from NSC.

### Box 2: Equivalence of NMF to STDPH and neural mechanisms of NSC

Spike-timing dependent plasticity (STDP) is a form of Hebbian learning in which synaptic weight changes depend on the relative timing between pre- and post-synaptic spikes [59, 60]. If pre-synaptic spike counts integrated over a critical window precede those of the post-synaptic neuron, then long-term potentiation (LTP) is induced, strengthening the weight. Otherwise, long-term depression (LTD) is induced, suppressing the weight. STDP is ubiquitous throughout the brain across the lifespan and can take on different forms [68, 69]. Simulations of STDP can result in runaway feedforward excitation such that produces unrealistic firing rates. To obviate this problem in biology, the brain uses homeostatic mechanisms that modulate synaptic inputs and neuronal firing thresholds [70].

We focus on synaptic scaling, a particular form of homeostasis which multiplicatively scales weights up or down depending on the average firing rate of the postsynaptic neuron, which has been demonstrated to stabilize STDP [20, 71, 72]. We refer to this combination of STDP and homeostatic synaptic scaling as STDPH.

We have shown that, given a network with fewer output than input neurons and full connectivity from the input layer to the output layer, STDPH iteratively acts to preserve the information in the first layer with the output layer neurons. Given an input layer of neurons (represented by matrix **V**) connected to a single output layer neuron (represented by row vector 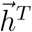) we have proven that the STDPH rule iteratively updates the synaptic weights in 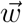 in such a way as to minimize the reconstruction error of 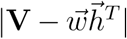, and that the NMF update rule is mathematically equivalent to the STDPH update rule [20].

To induce sparsity in networks of neurons, there must be a mechanism that causes each output layer to learn a different component of the total information represented by the input layer [73], which may be achieved by competition among units. Lateral or feedforward inhibition is often used to induce competition in SNNs, since it offers a biologically plausible mechanism to implement a winner-take-all (WTA) architecture [64], although other groups have suggested that thresholding can also achieve sparse codes [63]. Therefore, NSC can be attained in a two-layer network with STDPH and a WTA architecture via inhibitory connections within or between layers. Interestingly, popular implementations of NMF (e.g., MATLAB, Scikit-Learn) include a constraint on the L1 norm of **H** that automatically lead to sparsity. Alternatively, an explicit sparsity level can be incorporated into models of NMF, such as in Hoyer’s model [12].

### Box 3: Evolutionary Algorithm and SNN Evaluation

Evolutionary algorithms are a class of optimization algorithms loosely based on concepts from evolution. A set of individuals (in this case, spiking neural networks (SNNs)) have a set of parameters to be evolved, which are initialized for the first generation with size *N*. The population undergoes a training and a testing phase followed by evaluation as determined through a user-defined function that measures how well individuals in the population meet the desired criteria. The resulting value is the individual’s level of *fitness*. A certain number of individuals are then chosen as parents for the next generation, which can undergo *crossover* (i.e., where a child inherits parameters from its parents whose values have been swapped at random) and/or *mutation* (i.e., where a child inherits parameters from its parents with values randomly chosen from a distribution of values) to produce a new generation of individuals. This process continues until the fitness of the population no longer improves.

Carlson and colleagues [74] developed an automated tuning framework that incorporates a plugin to CARLsim [75] for running evolutionary algorithms using the ECJ library [76]. The plugin is used to pass information about the SNN population (e.g., number of generations to run, parameters to be evolved and their ranges, mutation and crossover rates) back and forth between CARLsim and ECJ (Fig. 4). The networks in the population are created, trained, tested, and evaluated using CARLsim. The resulting fitness scores for each individual in the population are then handed off to ECJ, which determines the parameters for the next generation according to the selection criteria defined in the parameter file, and the cycle repeats.

**Figure 4:**
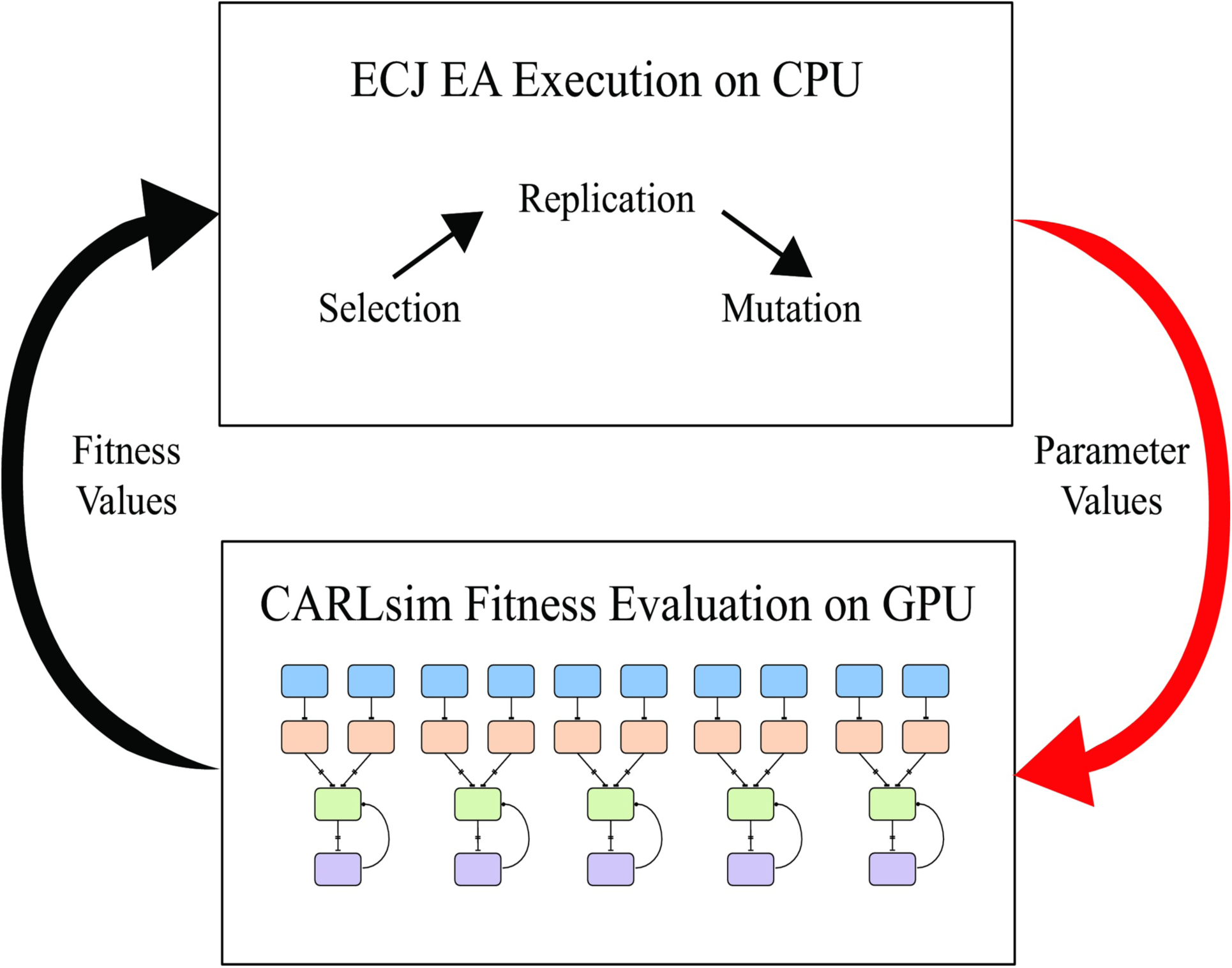
CARLsim-ECJ pipeline for evolving SNNs using ECJ (adapted from [74]). ECJ is used to initialize the parameters being tuned, which are passed to CARLsim and used to set the parameters of each network in the population of SNNs (red arrow). The SNNs are initialized, run, and evaluated using CARLsim. Following fitness evaluation using some user-defined criteria, the resulting fitness values for each individual in the population are sent to ECJ (black arrow). ECJ chooses parents from the population and performs selection, replication, and mutation on the parameters of the parent individuals in order to initialize the next generation of SNNs. This continues for a user-defined number of runs or until convergence.

In the experiments in which SNNs were used to replicate neurophysiological data, the evolutionary algorithm was used to optimize STDPH parameters in the model, corresponding to the temporal window over which spike times were integrated, the amount each synaptic weight increased or decreased, the initial weight range of each group of synaptic connections, and the target baseline firing rates of excitatory and inhibitory neurons. During each generation of the evolutionary run, models were trained on trials randomly selected from half of the available dataset, then tested on trials randomly selected from the latter half reserved for testing. At the end of each generation, synthetic neural activity was correlated with electrophysiologically recorded neural activity, which served as the fitness metric for the algorithm. After the model finished an evolutionary run, SNN activity closely matched the electrophysiological activity.

## Glossary

Basis functions: A lower-dimensional set of linearly independent elements that can represent a high-dimensional input space given a weighted sum of these elements, where the weight of each element is defined by a separate hidden component.
Dimensionality reduction: The process of reducing the dimensionality of a space to the lowest possible space that encapsulates the variance of the original data via feature extraction. In the case of neuronal firing rate patterns, this means representing all possible firing rate patterns in the brain region using the smallest possible subset of the neurons.
Factor analysis: A statistical method used to describe variability among observed, correlated variables in terms of a potentially lower number of unobserved, uncorrelated variables called factors (or latent variables).
Receptive field: The structure and boundaries of an individual neuron’s pattern of response to various kinds of incoming stimuli.
Spike-timing dependent plasticity: A Hebbian-inspired learning rule in which weight updates are computed based on the precise spike times of pre- and post-synaptic neurons that induce either long term potentiation or long term depression in the synapse, depending on whether the total pre-synaptic spike count preceded the total post-synaptic spike count, integrated over a critical temporal window.

